# A reconstituted mammalian APC-kinesin complex selectively transports defined packages of mRNAs

**DOI:** 10.1101/701250

**Authors:** Sebastian Baumann, Artem Komissarov, Maria Gili, Verena Ruprecht, Stefan Wieser, Sebastian P. Maurer

## Abstract

Through the asymmetric distribution of mRNAs cells spatially regulate gene expression to create cytoplasmic domains with specialized functions. In mammalian neurons, mRNA localization is required for essential processes as cell polarization, migration and synaptic plasticity underlying long-term memory formation. The essential components driving cytoplasmic mRNA transport in neurons and mammalian cells are not known. Here, we report the first reconstitution of a mammalian mRNA transport system revealing that the tumour suppressor adenomatous polyposis coli (APC) forms stable complexes with the axonally localised β-actin and β2B-tubulin mRNAs which are linked to a heterotrimeric kinesin-2 via the cargo adaptor KAP3. APC activates kinesin-2 and both proteins are sufficient to drive specific transport of defined mRNA packages. Guanine-rich sequences located in 3’UTRs of axonal mRNAs increase transport efficiency and balance the access of different mRNAs to the transport system. Our findings reveal for the first time a minimal set of proteins capable of driving kinesin-based, mammalian mRNA transport.

## Introduction

By localising mRNAs and producing proteins locally, cells spatially control gene expression, allowing them to build local protein networks with specialized functions ^1–3^. Different cell types localise up to thousands of different mRNAs to specific domains which is required for a wide range of cellular functions ^4–7^. New imaging techniques allowed to visualise individual-mRNA transport events in living neurons ^8,9^ and reconstitutions revealed the first essential components of yeast and drosophila mRNA transport systems^10–13^. Still, as many building blocks of these reconstituted systems lack mammalian homologues, it remained unknown how any mammalian mRNA is loaded onto motor proteins, how copy number of transported mRNAs is controlled and what molecular mechanisms generate specificity to transport only certain mRNAs. Provided the essential building blocks are identified and can be purified, an in vitro reconstitution could provide answers to these questions for the first time.

Adenomatous polyposis coli (APC) binds the KAP3 cargo-loading subunit of the kinesin-2 KIF3A/B/KAP3 ^14^, microtubules ^15,16^ and guanine-rich RNA sequences present in 3’UTRs of axonally localised mRNAs ^17^. The localisation of APC to axonal growth cones depends on KIF3A/B/KAP3 ^18^ and APC, in turn, is required for the axonal localisation and translational regulation of several mRNAs ^17^. While this suggests that APC might link axonally localised mRNAs to the microtubule-based transport machinery, direct evidence for this hypothesis has been missing. To address this, we purified ^19,20^ fluorescently tagged ^21,22^ full-length mouse APC, the KIF3A/B/KAP3 heterotrimer (KIF3ABK) and obtained synthesised β-actin (βact_wt_) and β2B-tubulin (β2Btub_wt_) mRNA fragments containing APC binding sequences. We then reconstituted APC-mediated mRNA transport in vitro (movie S1) and used a TIRF-M (Total-Internal-Reflection-Microscopy) assay (Fig. S1a) with single-molecule sensitivity to directly visualise and dissect the machinery and regulatory mechanisms driving mammalian mRNA transport.

## Results

To test whether full-length APC could function as linker between RNAs and kinesins, we first measured the interaction strength between full-length APC-GFP (Fig. S1b) and a minimal β2Btub-mRNA 3’UTR fragment (β2Btub_wtmin_) containing a guanine-rich APC interaction motif ^17^ (Table S1) using Microscale Thermophoresis (MST, Fig. 1a). In a second experiment, we measured the interaction strength between APC-GFP and KAP3 (Fig. 1b, Fig. S1c). Both measured affinities are in the lower nanomolar range, making the assembly of a ternary RNA-transport complex a possible scenario. To further test whether these interactions are sufficient to drive processive mRNA transport, we next combined KIF3ABK (Fig. S1d), TMR-labelled APC and an Alexa647-labelled β2Btub-mRNA fragment (β2Btub_wt_, Table S1) in a TIRF-M in vitro motility assay (movie S2). The components assembled into processively moving complexes containing both APC and β2Btub_wt_ (Fig. 1c).

**Fig. 1.**
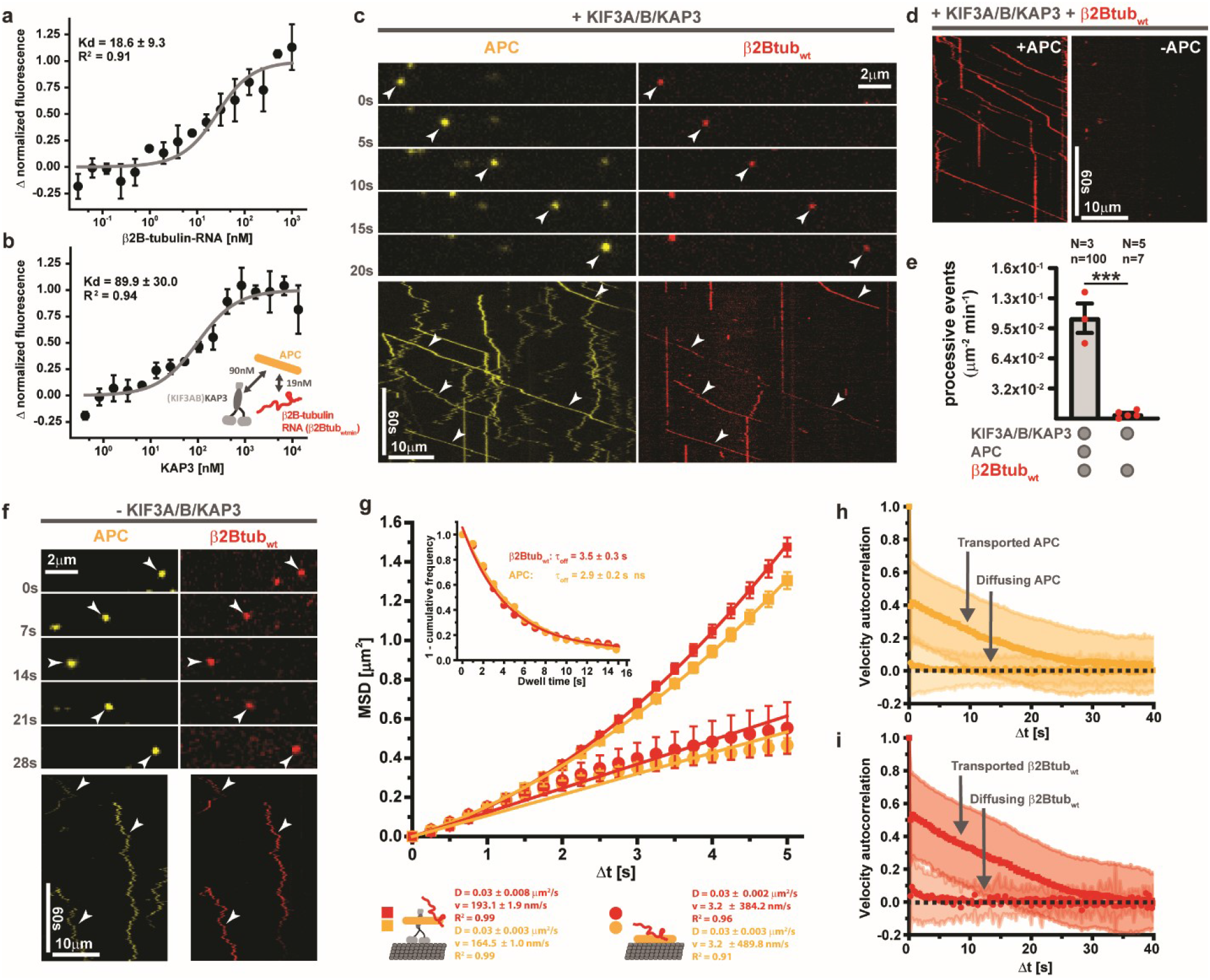
APC and kinesin-2 are necessary and sufficient to transport β2Btubulin-mRNA. **(A & B)** Full-length APC binds KAP3 and β2Btubulin-RNA (β2Btub_wtmin_) with different affinities. Interactions of APC-GFP with β2Btub_wtmin_ (A) and KAP3 (B) were measured by Microscale Thermophoresis in solution. The inset in (B) shows a summary of affinities measured between RNA-transport complex components. Error bars: SD. **(C)** KIF3ABK transports RNPs containing APC-TMR and Alexa647-β2Btub_wt_. Upper panel: time sequence from a TIRF-M assay containing 750 pM KIF3ABK, 80 pM APC-TMR and 2 nM Alexa647-β2Btub_wt_. Lower panel: A kymograph showing multiple run events of APC-TMR and Alexa647-β2Btub_wt_ complexes, indicated by white arrowheads. **(D)** APC is essential for β2Btub_wt_ transport. The kymographs show Alexa647-β2Btub_wt_ signals from TIRF-M experiments with (left) or without (right) APC. **(E)** Quantification of processive β2Btub_wt_ run events in the presence and absence of APC. N: number of independent experiments, n: total number of events. Error bars: SEM. Statistical significance was evaluated with an unpaired, two-tailed t-test. ***p<0.001. **(F)** APC recruits β2Btub_wt_ to the microtubule lattice in the absence of KIF3ABK. Upper panel: time sequence from a TIRF-M assay containing 40 pM APC-TMR and 2 nM Alexa647-β2Btub_wt_. Lower panel: Kymographs showing APC-RNA co-diffusion events (white arrowheads). **(G)** Mean-Squared-Displacement (MSD) plots of APC-TMR and Alexa647-β2Btub_wt_ from the experiments shown in (C) and (F). Error bars: SEM. Inset: the dwell times of APC-TMR and Alexa647-β2Btub_wt_ in the absence of KIF3ABK on the microtubule lattice. Statistical significance was evaluated with a Mann-Whitney test on the raw data ***p<0.001. **(H & I)** APC-β2Btub_wt_-complex lattice diffusion is not biased. Velocity autocorrelations of transported and lattice-diffusing APC-TMR (H) and Alexa647-β2Btub_wt_ (I) are shown. Error bars: SD.

To assess whether APC is essential for β2Btub_wt_ transport, we performed experiments with and without APC resulting in the loss of processive RNA movement in the absence of APC (Fig. 1d&e). Without the motor protein, APC (movie S3) and APC-β2Btub_wt_ ribonucleoprotein complexes (APC-RNPs, movie S4) bind and diffuse on the microtubule lattice (Fig. 1f,) showing that the reported microtubule binding ^15^ and RNA binding ^17^ activities of APC are not mutually exclusive at a saturating RNA concentration (Fig. S1f).

To dissect the different types of motility observed, we next used a single-particle tracking method ^23^ to create individual trajectories of diffusing and transported APC-TMR and Alexa647-β2Btub_wt_ molecules (Fig. S1g&h). To avoid bias, the entire field of view of entire TIRF-M movies was analysed. We then computed the mean-squared displacement ^24^ (MSD) of APC and β2Btub_wt_ signals in dual-colour experiments in the presence and absence of KIF3ABK (Fig. 1g) which shows that kinesin addition induces processive movement of APC and β2Btub_wt_. Dwell time measurements of diffusing APC-RNPs in the absence of KIF3ABK, over time frames in which no bleaching occurs (Fig. S1i&j), yield results of nonsignificant difference for APC and β2Btub_wt_, indicating that the APC-RNA interaction is more stable than the APC-microtubule interaction (Fig. 1g, inset). A velocity autocorrelation analysis (Fig. 1h&i) demonstrates that opposed to transported APC-RNPs lattice diffusing RNPs to do not show any directionality. These experiments reveal that APC couples β2Btub_wt_ with high affinity to a kinesin cargo adaptor and microtubules to enable both, directed and none-biased diffusive RNA transport.

Next, we assessed whether KAP3 which is thought to function as adaptor between APC and the KIF3AB heterodimer ^14^ (Fig. S2a) and which is required for axonal localisation of APC ^18^ is necessary for β2B-tubulin RNA transport. In the absence of KAP3, directed β2Btub_wt_ transport was not detectable, while adding back separately purified KAP3 (Fig. S1c) to the assay restored processive β2Btub_wt_ movement (Fig. 2a-c). Using Size-Exclusion-Coupled Multi-Angle-Light-Scattering (SEC-MALS), we confirmed that separately purified KAP3 can interact with KIF3A (Fig. S2c) dimers to form a trimeric complex (Fig. S2d). The mean velocity measured for processive (displacement > 4μm, Fig. S2e) β2Btub_wt_ transport complexes was 0.57 μm/s (Fig. 2d) which matches the reported velocity for KIF3ABK in cells ^25^ and in vitro ^26^ and also the reported velocities for axonally localised RNAs in cultured neurons ^9,27^.

**Fig. 2.**
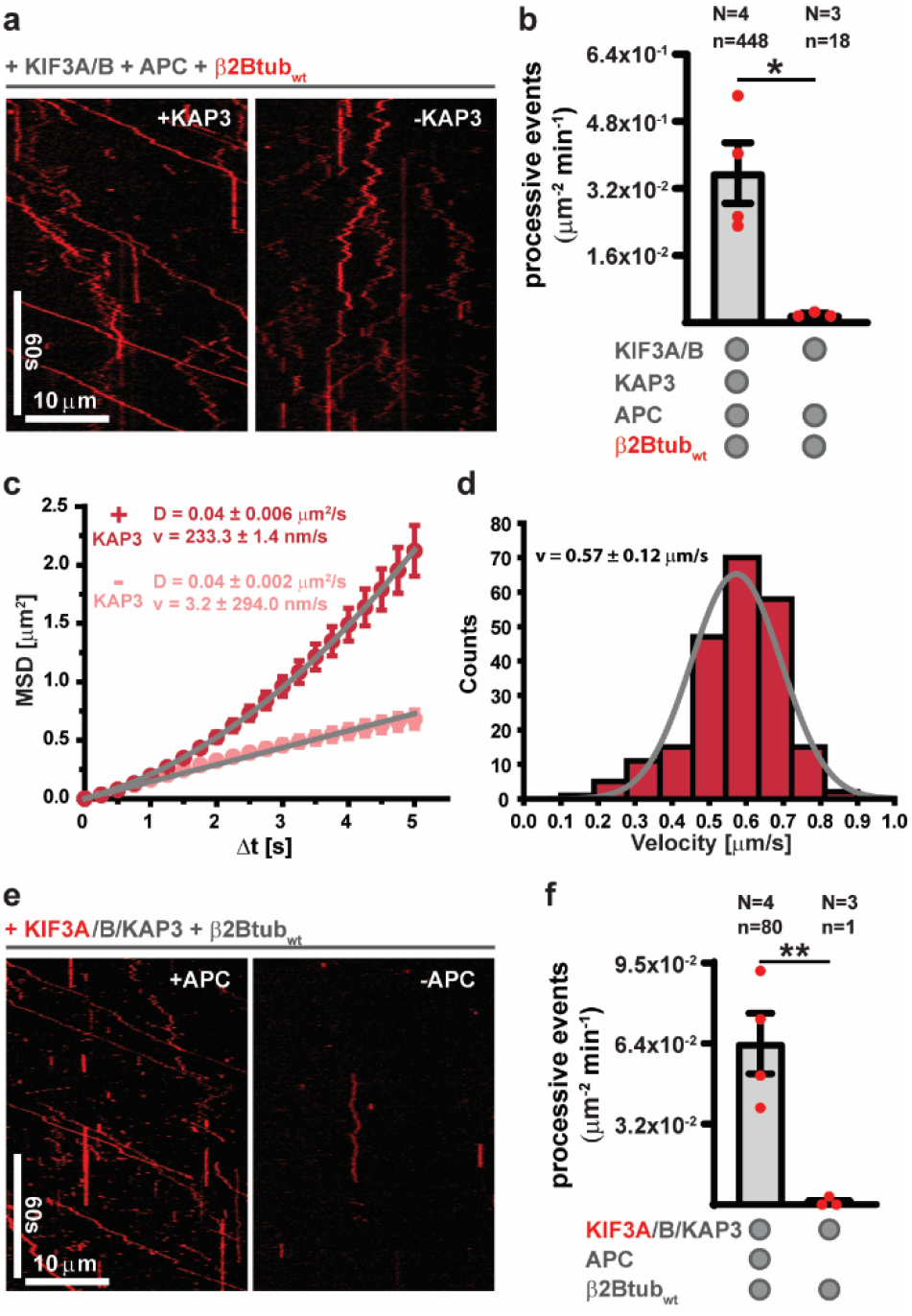
Function of the individual mRNA transport complex components. **(A)** KAP3 is required for processive RNA transport. The kymographs show Alexa647-β2Btub_wt_ signals from TIRF-M experiments containing 750 pM KIF3AB, 150 pM APC, 2 nM Alexa647-β2Btub_wt_ and 500 pM KAP3 (left) or no KAP3 (right). **(B)** Quantification of processive Alexa647-β2Btub_wt_ run events in the presence and absence of KAP3. Error bars: SEM. **(C)** MSD plots of Alexa647-β2Btub_wt_ motility (displacement >1.1 μm to include none-processive events in experiments lacking KAP3 from experiments shown in (A). Error bars: SEM. **(D)** Mean instantaneous velocity distribution of processive (displacement > 4 μm) Alexa647-β2Btub_wt_ complexes. Grey line: A gauss fit to velocity distribution. **(E)** APC recruits and activates KIF3ABK. The kymographs show Alexa647-KIF3ABK (750pM) signals from TIRF-M experiments containing 2 nM TMR-β2Btub_wt_ and 80 pM APC (left) or no APC (right). **(F)** Quantification of processive Alexa647-KIF3ABK run events in the presence and absence of APC. Error bars: SEM. In B and F statistical significance was evaluated with an unpaired, two-tailed t-test. **p<0.01, *p<0.05. N: number of independent experiments, n: total number of events.

Microtubule binding proteins as APC often function as catalysators for microtubule recruitment and processive movement of kinesins ^28,29^. APC, harbouring an N-terminally localised KAP3 binding site ^14^ and a C-terminally localised Tau-like microtubule binding site ^30^, could function as recruitment factor for KIF3ABK. To test this, we purified full-length, heterotrimeric, Alexa-647 labelled KIF3ABK (Fig. S1d) and tested its activity with in vitro motility assays in the presence and absence of APC (Fig. 2e&f). Addition of APC triggered microtubule recruitment and processive runs of Alexa647-KIF3ABK (movie S5), demonstrating that indeed APC functions as kinesin-2 activator in vitro.

In neurons, mRNAs are transported in defined packages containing only a few mRNA molecules ^8,9,31,32^. Interestingly, the quantity of mRNAs found in mRNA transport complexes is flexible within limits; while the majority of complexes contain one or two of the same mRNAs, a smaller fraction contains more ^8,9^. To analyse whether our reconstituted system recapitulates this key-property of neuronal mRNA transport complexes and to understand the architecture of our reconstituted transport complexes, we measured the numbers of motor, APC and RNA molecules in reconstituted β2B-tubulin RNA transport complexes at a saturating RNA concentration (Fig. S1f). As calibration standard to determine single-fluorophore intensities, we used immobilised SNAP-KIF3A (Fig. S3a&b) labelled with either Alexa647 or TMR and first extracted dual- and single-fluorophore intensities from bleaching steps (Fig. 3a&b). Bleaching step analysis provide a reliable reference for fluorophore brightness but cannot be used to measure the number of molecules in motile transport complexes. Hence, we next confirmed that intensity distributions generated by single-particle tracking of immobilised Alexa647-KIF3A dimers (Fig. 3c) match intensities obtained from bleaching step analysis.

**Fig. 3.**
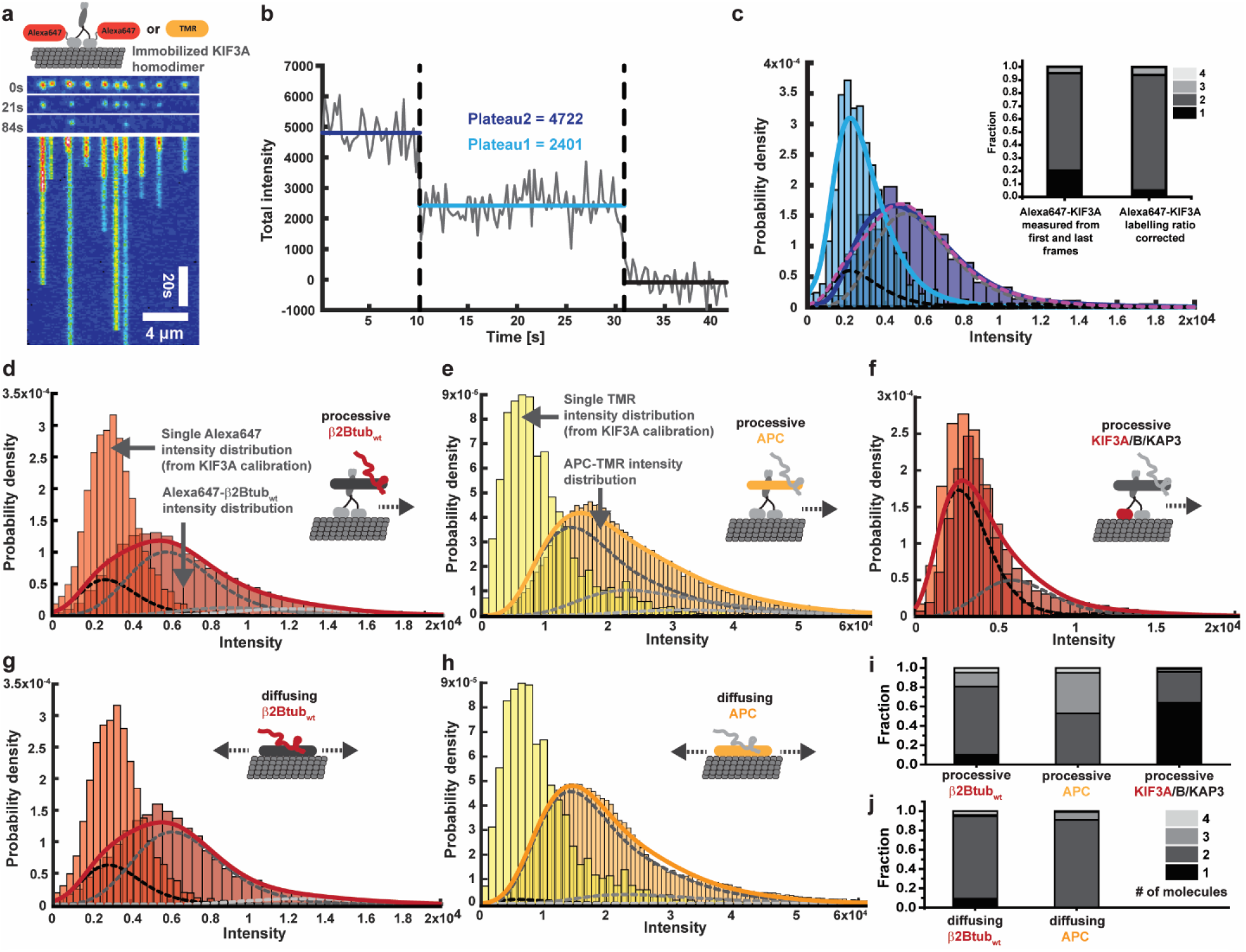
Fluorescence quantification of motile complexes reveals the mRNA transport complex architecture. **(A)** Bleaching of an Alexa647-KIF3A homodimer calibration sample. The cartoon on the top schematizes the experimental setup; 10 pM homodimeric SNAP-KIF3A labelled either with Alexa647 or TMR was imaged on paclitaxel-stabilized microtubules in the presence of AMPPNP to immobilize the kinesin. The central three images and kymograph at the bottom show an exemplary time course of Alexa647-KIF3A bleaching. **(B)** A bleaching trace obtained from a single Alexa647-KIF3A dimer indicating total intensities of one and two Alexa647 fluorophores. **(C)** The blue distributions show total intensities derived from single-particle tracking of immobilized KIF3A-Alexa647 dimers (light blue = derived from the first 10, dark blue = derived from the last 20 frames). Correspondingly, the light and dark blue lines indicate kernel-density-function (KDF) fits to the shown distributions. The dotted magenta line shows to the least square fit of the sum of the monomer and multimer fractions (black and grey dotted lines), derived from iterative convolutions of the monomer distribution (light blue line). The inset bar graphs show the measured fraction of KIF3A mono and multimers and the real fractions obtained after correcting for the labelling ratio of KIF3A (Table S2). **(D-F)** Quantification of molecules in individual RNA-transport complex components. Using single-particle tracking, the total intensities of KIF3ABK, APC and β2Btub_wt_ was measured in processive (displacement > 4 μm) complexes. For (D+E) experiments contained 750 pM KIF3ABK, 80 pM APC-TMR and 2 nM Alexa647-β2Btub_wt_. For (F), 750 pM Alexa647-KIF3ABK, 80 pM APC and 2 nM TMR-β2Btub_wt_. **(G + H)** Intensity distributions of microtubule lattice diffusing RNPs from assays containing 40 pM APC-TMR and 2 nM Alexa647-β2Btub_wt_. **(I)** Number of molecules of RNA-transport complex components derived from (D-F) after labelling ratio (Table S2) correction. **(J)** Number of molecules in diffusive RNPs derived from (G&H) after labelling ratio correction. The schematized complexes depicted in D-H illustrate the experimental setup and the transport complex component analysed (coloured). For D-H at least three independent experiments were analysed providing at least 75 tracks containing > 11.000 total intensities from detected particles. Light red or light-yellow distributions show intensity distributions of Alexa647- or TMR-Kif3A from calibration experiments. Coloured lines (red or yellow) indicate KDF. Dotted lines indicate monomer and multimer fractions greyscale according to legend in (J).

We then used single-particle tracking to measure the total intensities of processively moving Alexa647-β2Btub_wt_(movie S6), APC-TMR and SNAP-KIF3ABK (with only KIF3A being Alexa647 labelled) in experiments in which either the kinesin and RNA or APC and RNA were labelled (Fig. 3d-f). The same analysis was done for microtubule lattice-diffusing APC-RNPs in the absence of the kinesin (Fig. 3g&h). We find that on average a single KIF3ABK hetero-trimer transports one or two APC-dimers that carry RNAs at an equal stoichiometric ratio (Fig. 3I). Lattice diffusing APC is mainly a dimer, which is on average bound by one RNA per APC monomer under saturating conditions (Fig. 3j). These results demonstrate that APC and KIF3A/B/KAP3 assemble into RNA transport complexes showing the same confined variability in RNA-cargo quantity per transport complex as observed in cells.

mRNA transport machineries must be selective for a subset of the thousands of expressed mRNAs and at the same time balance transport frequencies of different mRNAs despite large differences in expression levels; also mRNAs present at low copy numbers must be transported. To first test whether our reconstituted system is selective, we conducted experiments with βact (βact_wt_) and β2Btub 3’UTR fragments which carry different variants of the APC binding motifs ^17^. In addition, we designed RNAs in which APC motifs are mutagenized to contain a low number of guanines. Comparing each wild type RNA to its mutated version shows that the APC-RNA transport system is selective (movie S7&8) giving clear preference to the wild type RNAs in both cases (Fig. 4a&b). At the same time these experiments show that APC can link different G-motif containing RNA sequence found in axonal mRNAs to kinesin-2. As β-actin mRNA is at least 10-fold more abundant than β2B-tubulin mRNA in cortical and hippocampal neurons ^33^, we next asked how an mRNA transport system compensates for different cargo concentrations in a situation of competition. At a 10-fold access of βact_wt_ over β2Btub_wt_, both RNAs are transported (Fig. 5a). Decreasing β2Btub_wt_ concentrations, while keeping the βact_wt_ concentration constant, increases the fraction of βact_wt_ bound to microtubules in an APC dependent manner (diffusive and processively transported, Fig. 5b). Analysing the total number processive βact_wt_ transport events showed that more βact_wt_ is transported with decreasing β2Btub_wt_ concentrations (Fig. 5c, movie S9). While the dwell times of both RNAs in the transport complexes do not show a significant difference (Fig. 5d), indicating that again RNA off rates are controlled by motor or APC off rates, the dwell time measured was twice as high compared to what we measured for lattice diffusing APC-RNPs (Fig. 1g, inset). This potentially stems from the additional microtubule attachment point KIF3ABK provides for APC. To test further what causes the observed preference of our reconstituted system for β2Btub_wt_ RNA, we conducted MST experiments with both RNAs and found that βact_wt_ had a 5-fold lower affinity to APC than β2Btub_wt_ (Fig. 5e). These results show that APC has different affinities to different mRNA localisation signals, which fine-tunes transport frequencies of different axonally localised mRNAs.

**Fig. 4.**
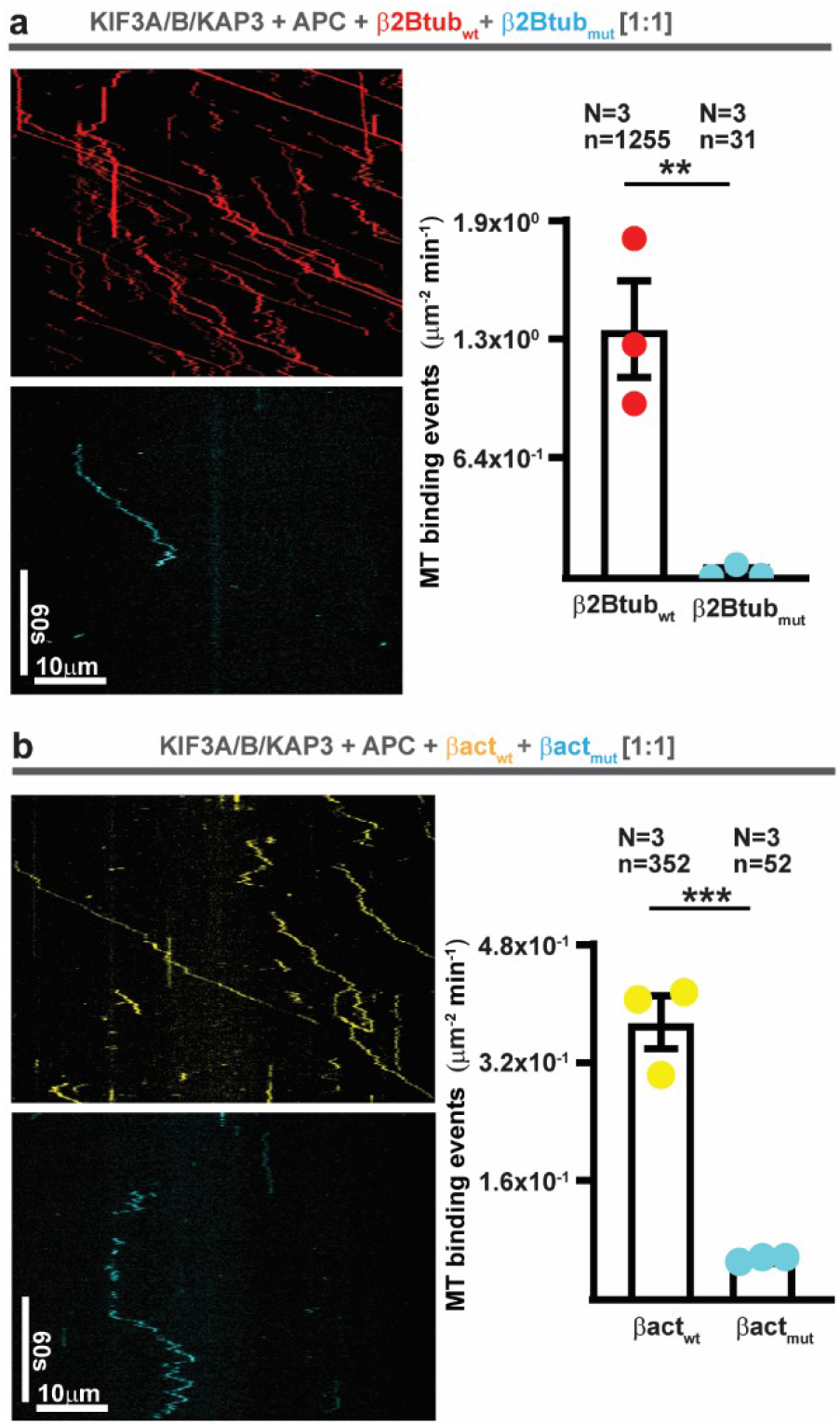
APC selectively linksβ-actin and β2B-tubulin mRNAs to kinesin-2. **(A)** Kymographs showing Alexa647-β2Btub_wt_ and TMR-β2Btub_mut_ in experiments containing 750 pM KIF3ABK and 150 pM APC. Equimolar RNA concentrations of 500 pM were used. Right panel: quantification of Alexa647-β2Btub_wt_ and TMR-β2Btub_mut_ microtubule binding events (diffusive & processive). **(B)** Same as in A, but comparing TMR-βactin_wt_ to Alexa647-βactin_mut_ RNA using equimolar RNA concentrations of 3000 pM. Right panel: quantification of TMR-βactin_wt_ and Alexa647-βactin_mut_ microtubule binding events (diffusive & processive). Error bars in (A+B): SEM.

**Fig. 5.**
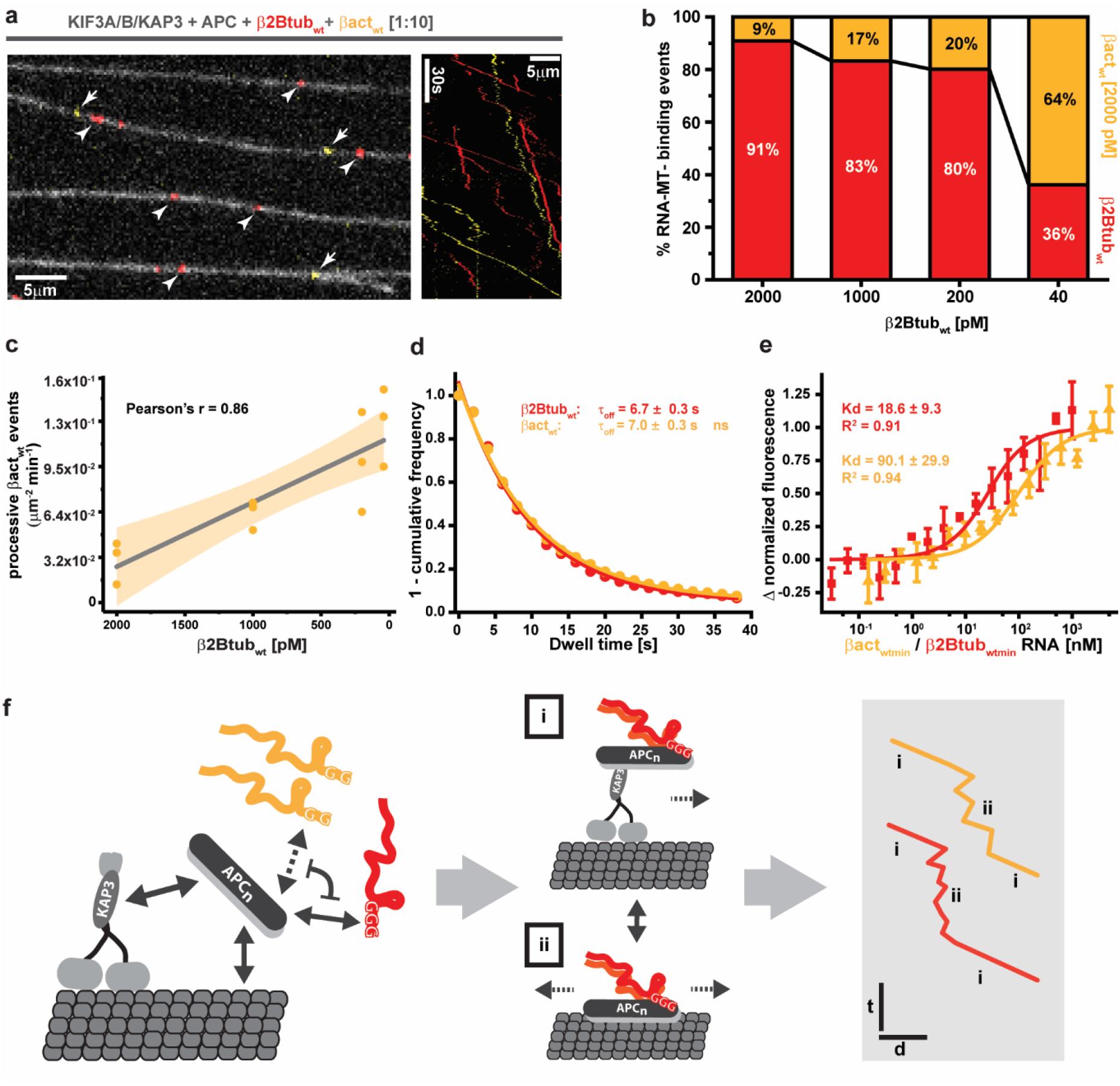
The affinity of APC to different mRNAs controls mRNA transport frequency. **(A)** An overview TIRF-M image (left) and kymograph (right) showing Alexa647-β2Btub_wt_ and TMR-βactin_wt_ transport in the same experiments at 1:10 molar ratio (200 pM:2 nM). Arrows and arrowheads point to TMR-βactin_wt_ and Alexa647-β2Btub_wt_ RNPs, respectively. **(B)** A titration of Alexa647-β2Btub_wt_ from 2000 to 40 pM leads to an increase of the relative amount of TMR-β2Btub_mut_ transport per experiment. **(C)** Plot of processive run events of TMR-βactin_wt_ (displacement > 4 μm) per experiment in dependence of the free Alexa647-β2Btub_wt_ concentration. Grey line: Linear regression fit. Yellow-shaded area: 95% confidence band. **(D)** Dwell time distribution of β2Btub_wt_ and βact_wt_ RNAs in experiments containing 750 pM KIF3ABK, 150 pM APC, 40 pM Alexa647-β2Btub_wt_ and 2000 pM TMR-βact_wt_. Coloured solid lines: monoexponential decay fits to dwell time distributions **(E)** MST measurement of the affinity of βactin_wt_ and β2Btub_wt_ RNA to APC-GFP. Error bars: SD. Solid lines: One-site binding model fit to the measured normalized fluorescence. **(F)** Model of transport RNP assembly. Oligomerising APC binds guanine-rich motifs located in mRNA 3’UTRs, microtubules and KAP3 with different domains. mRNAs with different G-motif variants compete for APC binding. APC-RNPs move by two different modes: **(i)** active transport by kinesin-2 **(ii)** microtubule lattice diffusion. In A+B statistical significance was evaluated with an unpaired, two-tailed t-test. ***p<0.001 **p<0.01. In F, statistical significance was evaluated with a Mann-Whitney test on the raw data. N: number of independent experiments, n: total number of events.

## Discussion

Our results show that APC selectively links RNAs with guanine-rich sequences found in 3’UTRs of mRNAs to the heterotrimeric kinesin-2 KIF3A/B/KAP3. APC’s microtubule, RNA and KAP3 binding activities recruit the heterotrimeric kinesin-2 to the microtubule and trigger the assembly of processive and diffusive mRNA transport complexes (Fig. 4H). We observe a mixture of diffusive and directed motion of APC-RNPs which was also observed for β-actin mRNA in neurons ^8,9,34^. Being able to switch from one transport mode to the other can be of advantage; diffusion is faster than directed transport over shorter distances, it is not affected by microtubule orientation and it allows switching of protofilament tracks on microtubules to navigate around roadblocks ^35^.

Our stoichiometric analysis of single reconstituted mRNA transport complexes matches the numbers of mRNAs found in neuronal mRNA transport complexes but detects a smaller fraction of single-RNA complexes. This could be because it is technically challenging to measure RNA labelling ratios in vitro and in cells. APC’s oligomerisation domains provide the means to pack different numbers of APC-RNA complexes on a kinesin, potentially providing a mechanism for cargo-concentration driven loading of transport complexes. At the same time cellular signalling events could modify the self-association abilities of APC ^36^ to trigger unloading of cargo-mRNAs. Indeed, the number of β-actin mRNAs per transport complex and density of RNAs decrease with increasing distance from the neuronal soma ^8^.

Our finding that different guanine-sequence variants present in β-actin and β2B-tubulin mRNAs lead to different transport efficiencies open an interesting conceptual framework for the evolution of mRNA transport signals; subtle sequence variations might balance the access of mRNAs to the transport system to guarantee transport also of low-abundant mRNAs. Transport motif-mediated fine tuning of mRNA on- and off-rates to adaptors as APC might be the molecular basis for the generation of different mRNA distributions observed in neurons ^4^.

The availability of functional full-length APC for in vitro studies now opens the door to understanding how APC integrates the signals it receives and the many activities it controls such as regulation of microtubule assembly ^37^, actin nucleation ^38^, micro-tubule-actin coordination ^39^, mRNA localization ^6^ and translation regulation ^17^. Identification of the first minimal, mammalian mRNA transport system will provide the means to reveal diversities in mRNA transport pathways in different cell types and to understand how mRNA transport, stability and translation are mechanistically connected.

## Acknowledgments

We thank the CRG BMS-PT core facility for support and training on instrumentation. We further thank Thomas Surrey, Vivek Malhotra, Ivo Telley and Christian Duellberg for critical comments on the manuscript.

## Funding

This work was funded by the Spanish Ministry of Economy and Competitiveness (MINECO) [BFU2017-85361-P], [BFU2014-54278-P], [BFU2015-62550-ERC], Juan de la Cierva-Incorporación Program [IJCI-2015-25994], the Human Frontiers in Science Program (HFSP) [RGY0083/2016] and the European Research Council (ERC) [H2020-MSCA-IF-2014-659271.]. We further acknowledge support of the Spanish Ministry of Economy and Competitiveness, ‘Centro de Excelencia Severo Ochoa’ [SEV-2012-0208].

## Author contributions

SB, SPM conceived the project. SB, MGS, AK, SPM performed experiments. SB, VR, SW, SPM analysed the data. SB, SPM wrote the article. SPM supervised the project.

## Competing interests

None.

## Data and materials availability

All data are available in the manuscript or the supplementary materials.

## Supplementary Materials

Materials and Methods

Fig.s S1-S3

Tables S1-S2

Movies S1-S9

## Materials and Methods

### Materials

Chemicals were obtained from Sigma if not stated otherwise. Single-fluorophore, 5’-end-labelled RNA fragments from mouse β-actin mRNA (accession number: NM_007393.5) and mouse β2B-tubulin-mRNA (accession number: NM_023716.2) were purchased from IBA Lifesciences (Germany).

TIRF microscopy experiments were performed on a iMIC (TILL Photonics, Germany) total internal reflection fluorescence (TIRF) microscope equipped with three Evolve 512 EMCCD cameras (Photometrics, UK), a 100x 1.49 NA objective lens (Olympus, Japan), a quadband filter (405/488/561/64, Semrock, USA) and four different laser lines (405 nm, 488 nm, 561 nm, 639 nm). An Olympus tube lens adds a post magnification of x1.33 which results in a final pixel size of 120.3 nm.

### Methods

#### Cloning and recombinant protein expression

Sequences encoding APC, KAP3, KIFA and KIFB proteins (*Mus musculus* APC, accession number: AAB59632, *Mus musculus* KAP3A, accession number: BAA08901.1, *Mus musculus* KIF3A, accession number: NP_032469.2 and *Mus musculus* KIF3B, accession number: NP_032470.3) were synthesized commercially and codon-optimized for expression in insect cells (ThermoFisher Scientific). PCR-amplified APC was inserted by Gibson assembly into a pCoofy27 ^19^ backbone for N-terminal fusion of 10xHis-ZZ-TEV and if required, C-terminal fusion of mGFP ^22^ or SNAP tag ^22^. A similar strategy was applied to obtain 9xHis-ZZ-SUMO-KAP3 and 9xHis-ZZ-SUMO-KIF3A, which were inserted into pCoofy17a ^19^. For expression of heterodimeric KIFA/B and heterotrimeric KIF3A/B/KAP3 we used the biGBac system ^20^. KIF3A was N-terminally fused to 6xHis-ZZ-HRV3C and C-terminally to a TEV cleavage site followed by a linker sequence (ILGAPSGGGATAGAGGAGGPAGLIN) and SNAP tag. KIF3B was fused to the N-terminal tags OneStrep-ZZ-HRV3C-mGFP-GGGS_linker-TEV. All constructs were validated by sequencing of the entire open-reading frame. Proteins were cloned downstream of the *polh* promotor into pLIB vectors and either KIF3A and KIF3B or KIF3A, B and KAP3 were combined by Cre/loxP recombination ^20^. Following simultaneous protein expression, cells were frozen in liquid N_2_ and stored at −80°C.

#### Protein biochemistry

##### APC purification

Cold APC purification buffer (100 mM NaPi, Sigma#S3139 and Sigma#S9390, 300 mM KCl, Sigma#P9333, 5 mM MgCl2 × 6 H_2_O, Sigma#M2670, 0.001% Brij35 ThermoFisher, #28316, 2.5 mM DTT, Sigma#D0632, 2.5 mM EDTA, Sigma#EDS) supplemented with protease inhibitors (Roche, #5056489001) and DNaseI (Roche, #10104159001) was added to frozen SF21 cell pellets expressing an APC construct. The pellet was thawed in a room temperature water bath and resuspended. The lysate was clarified by centrifugation (184.000 x g, 45 min, 4°C) and the supernatant was applied to a 5ml HiTrap IgG Sepharose FF column (GE Healthcare, #28-9083-66). After washing with APC purification buffer, GST-TEV protease was added to the column and the column was left at 4°C over night to cleave APC off its column-bound affinity and solubility tags. The next day, APC was eluted with APC purification buffer and labelled with SNAP-reactive dye (NEB#S9105S). After SNAP-labelling, APC was concentrated using Vivaspin concentrators (Sartorius), ultra-centrifuged (280.000 x g, 10 min, 4°C) and gel-filtered using a Superose6 10/300 GL column (GE Healthcare). Peak fractions were pooled, and the labelling ratio was determined using Nanodrop (ThermoFisher). The protein was aliquoted on ice and snap frozen in liquid nitrogen.

##### KIF3A/B(/KAP3) purification

Cold KIF3A/B purification buffer (80 mM NaPi, 200 mM KCl, 5 mM MgCl2 × 6 H_2_O, 0.001% Brij35, 2.5 mM DTT, 2.5 mM EDTA, 0.1 mM ATP, #A2383) supplemented with protease inhibitors and DNaseI was added to frozen SF21 cell pellets expressing a KIF3A/B construct. Thawing and lysate clearing was done as described for APC purification above. Next, the supernatant was applied to a StrepTrap column (GE Healthcare, 28-9075-48). The column was washed with KIF3A/B purification buffer and the protein was eluted in KIF3A/B purification buffer supplemented with 15 mM d-desthiobiotin (Sigma, D1411-1G). Depending on whether unlabelled or fluorescently tagged protein was needed, the eluate was incubated either with TEV and 3C or only 3C protease over night at 4°C. After SNAP labelling, the protein was concentrated and ultra-centrifuged as described above, and gel-filtered using a GE Superdex200 10/300 column. Peak fractions were pooled, the labelling ratio was determined using Nanodrop. The protein was aliquoted on ice and snap frozen in liquid nitrogen.

##### KIF3A purification

Cold KIF3A purification buffer (50 mM NaPi, 400 mM KCl, 2 mM MgCl2 × 6 H_2_O, 0.001% Brij35, 0.75 mM β-mercaptoethanol, Sigma#M3148, 0.2 mM ATP) supplemented with protease inhibitors and DNaseI was added to frozen BL21-RIL cell pellets expressing a KIF3A construct. Pellets were thawed and sonicated. The lysate was clarified by centrifugation (184.000 x g, 45 min, 4°C) and the supernatant was applied to a 5 ml HiTrap column (GE Healthcare) loaded with cobalt(II)-chloride hexahydrate (Sigma#255599). The column was washed with KIF3A purification buffer and the protein was eluted in KIF3A purification buffer supplemented with 500 mM imidazole (Sigma#I2399). The eluate was concentrated with a Vivaspin concentrator (Sartorius) and supplemented with 8 mM β-mercaptoethanol. Affinity tags were cleaved over night at 4°C using SUMO protease. In case of KIF3A-SNAP, the protein was concentrated after labelling using a Vivaspin concentrator (Sartorius), ultra-centrifuged (280.000 x g, 15 min, 4°C) and gel-filtered using a GE Superdex200 10/300 GL column. Peak fractions were pooled, and the labeling ratio was determined using Nanodrop. The protein was aliquoted on ice and snap frozen in liquid nitrogen.

##### KAP3 purification

Cold KAP3A purification buffer (50 mM HEPES, Sigma#H3375, 400 mM KCl, 2 mM MgCl2, 0.002% Brij35, 1 mM β-mercaptoethanol, 1 mM imidazole, 50 mM L-Arginine, Sigma#A5006, 50 mM L-Glutamate, Sigma#49449, pH 7.25) supplemented with protease inhibitors and DNaseI was added to frozen BL21-AI cell pellets expressing a KAP3A construct. Pellets were thawed and cells lysed by French press at 10.000 psi. The lysate was clarified by centrifugation (184.000 x g, 45 min, 4°C) and the supernatant was applied to a 5 ml HiTrap column (GE Healthcare) loaded with cobalt(II)-chloride hexahydrate (Sigma). The column was washed with KAP3A purification and buffer containing 10 mM Imidazole and the protein was eluted in KAP3A purification buffer supplemented with 400 mM imidazole. The eluate was concentrated with a Vivaspin concentrator (Sartorius) and supplemented with 8mM β-mercaptoethanol. Affinity tags were cleaved over night at 4°C using SUMO protease. KAP3A was concentrated using a Vivaspin concentrator (Sartorius), ultra-centrifuged (280.000 x g, 15 min, 4°C) and gel-filtered using a GE Superdex200 10/300 GL column. Peak fractions were pooled, and the protein was aliquoted on ice and snap-frozen in liquid nitrogen.

#### SNAP labelling of proteins

SNAP labelling was carried out at 15°C for 90 minutes using 3-fold access of dye over protein before gel filtration. Fresh DTT (2 mM) was added directly before labelling. Unbound dye was immediately removed using Zeba™ Spin desalting columns (Thermo Fisher).

#### SDS PAGE and western blotting

SDS-PAGE was performed using Mini-Protean system and TGX precast gels (Biorad). Proteins were stained with BlueSafe solution (NZYtech). Western blotting on SDS gels was performed with an Invitrogen iBlot2 device and fitting iBlot2 NC stacks (Invitrogen). The antibody for KAP3 detection was purchased from Santa Cruz Biotechnology (sc-55598 HRP).

#### Reproducibility

At least three independent experiments were performed for all tested conditions.

#### Microscale thermophoresis (MST) measurements

MST experiments were carried out with a Monolith NT.115 system (Nanotemper, Germany). 15 nM of APC-GFP was titrated on ice with increasing concentrations of KAP3 (13450 nM - 0.41 nM, β-act_wtmin_ (5000 nM – 0.15 nM) or β2Btub_wtmin_ (1000 nM – 0.03 nM) in final assay buffer: 25 mM Tris pH 7.4, 75 mM NaCl, 5 mM MgCl2, 0.025% Tween-20. RNA assay buffer was supplemented with 0.125 u SuperaseIn (Invitrogen) and 2.5 mM EDTA. 16 PCR tubes were filled with a 1:2 dilution of the titrants and a constant amount of APC-GFP was added. Tubes were shifted to RT and mixtures were transferred to 16 standard capillaries (Nanotemper, Germany) according to the manufacturer’s manual, which were allowed to equilibrate for 4 min at 30°C before measurement. Data were acquired using 40% MST and 50% LED (blue filter) power. Binding curves were obtained by plotting the fluorescent change 5 s after applying the MST gradient against the concentration of the titrated protein or RNA. Data from three replicates were averaged and fitted with a Kd model derived from the law of mass action ^40^:

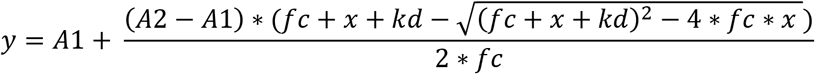

The concentration of the fluorescent molecule (fc, APC-GFP in our case) is constant at 15 nM while x corresponds to the concentration of the titrated ligand. A1 and A2 correspond to the normalized fluorescence value of the unbound and bound state, respectively.

#### Size-Exclusion Chromatography coupled Multi-Angle-Light Scattering (SEC-MALS) measurements

Storage buffer of KIF3A and KAP3 was exchanged against SEC-MALS buffer (30 mM HEPES, 200 mM KCl, 2 mM MgCl_2_, 10 mM β-mercaptoethanol, 0.05% NaN_3_) with Zeba™ Spin desalting columns (Thermo Fisher). Proteins were resolved either separately or after mixing KIF3A with KAP3A in a 2:1 molar ratio on a Shodex KW-804 column (Shodex, Germany). Following SEC fractionation, eluted protein was detected on a Wyatt Dawn 8^+^ eight angle light scattering instrument coupled to a Wyatt Optilab T-rEX online refractive index detector in a standard SEC-MALS format.

#### Polymerisation and stabilisation of microtubules

Microtubules were polymerised from bovine tubulin with stochastic incorporation of ATTO390 (Attotech, Germany) and biotin (ThermoFisher, #20217) labelled tubulin. Mixes of 36 μM unlabelled tubulin, 14 μM ATTO390-tubulin, and 10 μM biotin-tubulin were incubated in BRB80 (80 mM PIPES pH 6.85, 2 mM MgCl2, 1 mM EGTA) with 4 mM GTP (Sigma) for 25 min at 37°C in a total volume of 25 ul. Next, prewarmed BRB80 supplemented with 20 μM paclitaxel (Sigma) was added and incubation continued for 45 min at 37°C. Polymerised microtubules were pelleted at 20.000 x g for 5 min and resuspended in 50 ul BRB80 supplemented with 20 μM paclitaxel.

#### Motility chamber preparation

Glass surfaces were prepared as described previously ^41^. Motility chambers with a volume of ~10 μl were assembled by adhering glass cover slips functionalised with PEG and Biotin-PEG (Microsurfaces Inc, USA, #Bio_01(2007134-01)) to glass slides passivated with PLL(20)-g[3.5]- PEG(2) (SuSos AG, Duebendorf, Switzerland) using 2 parallel segments of double-sided adhesive tape. Chamber surfaces were passivated for 5 min on ice with 5% (w/v) Pluronic F-127 #P2443) in TIRF-M assay buffer (90 mM HEPES, 10 mM Pipes #P6757, 2.5 mM MgCl_2_, 1.5 mM EGTA #03777, 15 mM β-mercaptoethanol, 1.2 g KOH #S8045) supplemented with 50 ug/ml kappa-casein #C0406. Chambers were then flushed with 25 ug/ml neutravidin (Invitrogen, #A-2666) in assay buffer containing 200 ug/ml kappa-casein and instantly rinsed with assay buffer followed by warm up to RT. After 3 min microtubules were diluted 1:15 in RT assay buffer and added to the chamber followed by 3 min incubation.

#### In vitro motility assay

Constituents of motility assays were incubated on ice for 15 min in TIRF-M assay buffer supplemented with 4% PEG-3350 #1546547, 2.5 mM ATP and 0.5 U/μl SuperaseIn (Invitrogen, AM2694). Typical protein concentrations were: 10.5 nM APC, 52.5 nM KIF3ABK, 140 nM RNA. The preincubation mix was diluted 69.89-fold into final motility buffer lacking RNase inhibitor and containing additionally 0.32 mg/ml glucose oxidase (AMSBIO, Germany, #22778.02), 0.275 mg/ml Catalase #C40, 50 mM glucose #G7021, 50 ug/ml beta-casein #C6905 and 0.12% methylcellulose #M0512. Exact final protein and RNA concentrations are given in Fig. legends. Diluted mixes were applied to immobilised microtubules in the motility chamber for imaging at room temperature (30 ± 1°C).

#### TIRF microscopy

For triple colour imaging two channels were recorded alternating, either with two or four frames per second (250 ms or 125 ms exposure per channel) by switching laser lines 561 nm and 639 nm. ATTO390 microtubules were recorded as a single image after time lapse movies had been finalized. This protocol was chosen since ATTO390 bleaches comparably fast and total illumination time per cycle is saved to increase the frame rate in the other channels. For quadruple colour imaging 488 and 639 nm laser lines were switched on simultaneously and alternated with the 561 nm laser line, followed by a single image of ATTO390 MTs at the end. Laser powers as well as exposure times and acquisition frame rates were kept constant within a set of experiments and corresponding calibration samples to allow for direct comparisons between different conditions. For channel alignments, images with 100 μm TetraSpeck fluorescent beads (Invitrogen, UK) were recorded in all channels before experiments were started.

#### Analysis of TIRF-M data

For analysis of dynamic properties and the stoichiometry of APC-RNA complexes (APC-RNPs), TIRFM-movies were loaded into the FIJI software ^42^ and analysed using Trackmate ^23^. In all cases entire TIRF movies including the entire field of view were analysed to guarantee an unbiased analysis. Single particles were detected using the LoG (Laplacian of Gaussian) detector with an estimated spot diameter of 900 nm. Intensity thresholds for spot detection depended on the amount of fluorescent species used in the respective experiments and was kept constant for all comparisons of experiments using the same concentrations of fluorescent proteins or RNAs and the same imaging settings. To compute trajectories, we used the “Simple LAP tracker” with the following settings: maximal linking distance = 500 nm, maximal gap-closing distance = 500 nm, maximal frame gap = 4. To read out intensities of stationary KIF3A-Alexa647 and KIF3A-TMR calibration probes used in Fig. 3, a track duration cut off of > 3s was used to exclude blinking events and transient, none-specific binding. For the analysis of kinesin-2, APC and RNA dynamics (Figs. 1G, 1G inset, 1H, 1I, 2C, 3G, 3H, 4A, 4B, 4D and 4F) trajectories with a displacement of > 1100 nm were used to analyse only diffusing and processive particles and exclude stationary events that likely arise due to unspecific interactions. As APC-dependent microtubule lattice diffusion never showed a displacement > 4000 nm (Fig. S2E), this value was used as cut off when only processive complexes had to be analysed in Figs. 1H&I, 2D, 3D-F, 4E and 4F inset. The background fluorescence of TIRF-M data was measured by calculating the mean intensity in a 28×18 pixel rectangular box at five different positions (upper left & right, centre, lower left & right) of the field of view at four different time points of the movie. The obtained mean background fluorescence was subtracted from total spot intensities obtained by Trackmate single-particle tracking. We further always analysed the entire field of views of entire TIRF-M movies in all cases to exclude bias.

#### Analysis of transport-RNP dynamics

Trajectories of APC, RNA and motors obtained from Trackmate processing of TIRF-M data were further analysed using the MSDanalyzer toolbox ^24^ for MATLAB to compute the Mean-Squared-Displacement (MSD) and the velocity autocorrelation. To compare RNP-movements with and without kinesin-2, we fitted the equation *MSD* = *v*^2^*t*^2^ + 2*Dt* assuming a mixed motion of single particles due to APC-induced microtubule lattice diffusion and directed transport. For MSD analysis, in all Fig.s a displacement cut off of > 1100 nm was used. To test for directed motion of APC-RNA complexes, the velocity auto correlation of diffusing (displacement cut off > 1100 nm) and transported (displacement cut off > 4000 nm) APC-RNPs were compared in the absence or presence of the kinesin, respectively. For random diffusion, the individual displacement vectors of a track are uncorrelated resulting in a V_corr_ value of zero. In the case of directed motion, subsequent displacement vectors are comparable, resulting in a V_corr_ value above zero. To analyse dwell times of APC and β2B-tubulin RNA, 1-cumlative frequency distributions of the respective dwell times (=track duration) were fitted to the monoexponential decay function *y* = *y*_0_ + *Ae*^−*x*/*t*^. The decay constant *τ* was derived by *τ* = *t* ∗ ln(2). The mean velocity of RNA transport complexes was determined by computing the average of 224 individual mean track velocities. The mean track velocity is the mean of the instantaneous velocities of a track.

#### Measurement of single-fluorophore intensities on immobilized samples

To obtain intensities of single and multiple Alexa647 and TMR dyes under conditions comparable to RNA transport assays, we used homodimeric KIF3A-SNAP labelled with either Alexa647 or TMR. The motor was washed into TIRF-M assay chambers in imaging buffer containing 5 mM AMPPNP to induce strong coupling to the immobilized paclitaxel-stabilized microtubules. To allow bleaching the amount of glucose in the imaging buffer was reduced from 50 to 1 mM. Then movies were recorded using identical imaging settings as used for RNA transport experiments until most of the fluorophores were bleached. Intensities were obtained from the first 10 frames (multimeric signal) or last 20 frames (monomeric = single fluorophore signal) using total intensity counts obtained from Trackmate as described above. To detect single step photobleaching events the Fiji plugin TimeSeries V3_0 analyzer was used and intensity time tracks were processed in MATLAB using a custom-written changepoint detection algorithm to identify intensity plateaus indicative for monomeric or multimeric signals.

#### Determination of RNA transport complex stoichiometries

To measure the intensity distribution of lattice-diffusing APC-RNA complexes and APC-RNA transport complexes, four different experiments were conducted. For lattice diffusing APC-RNA complexes, 40 pM APC-GFP and 2000 pM Alexa647-β2Btub_wt_ were used.. To measure the stoichiometry of processive RNA-transport complexes, protein concentrations in the TIRF-M assays were as follows: for KIF3A/B/KAP3 intensity measurements, 750 pM Alexa647-KIF3A/B/KAP3, 80 pM APC and 2000 nM TMR-β2Btub_wt_ were used. For APC intensity measurements, 750 pM KIF3/A/B/KAP3, 80 pM APC-TMR and 2000 nM Alexa647-β2Btub_wt_. For RNA intensity measurements, 750 pM KIF3/A/B/KAP3, 80 pM APC and 2000 nM Alexa647-β2Btub_wt_. Imaging was done with triple-colour mode at four frames per second (described above). Kinesin, APC and RNA intensities were then obtained from Trackmate analysis and background intensities were subtracted as described above. Using a custom written MATLAB script, we calculated the probability density function (kernel density estimation) for both the monomeric intensity distribution (obtained from Alexa647-KIF3A or TMR-KIF3A bleaching experiments, pdf_mono_) and the intensity distributions of RNA-transport complex components (pdf_RNP_). In general, intensity distributions of Alexa647 and TMR fluorescent molecules are typically right-skewed and broader than Poissonian distributions with std~√N, N the number of photons in the signal. The pdf of the experimentally determined single-Alexa647 or -TMR intensity distribution pdf_mono_ was further used to calculate dimeric (pdf_di_) and higher multimeric (pdf_tri_, pdf_quat_) intensity distributions based on iterative convolutions of pdf_mono_ as described previously ^43^. To reveal the stoichiometry of the protein complex, a linear combination of pdf_mono_, pdf_di_, pdf_tri_ and pdf_quatt_ was fitted to individual pdf_RNP_ distributions using a least squares method, taking into account the given protein and RNA labelling ratios.

---

**Fig. S1.**
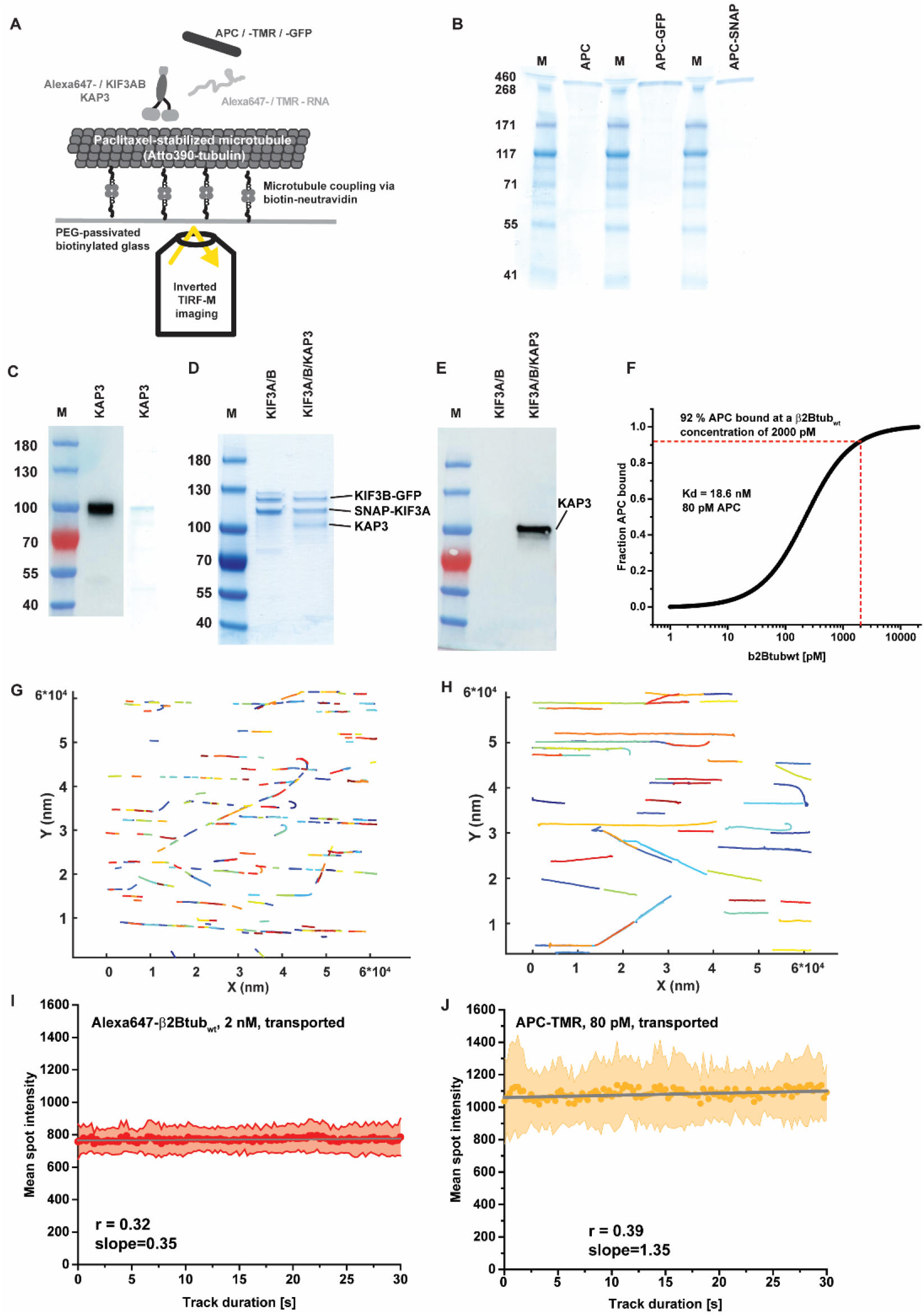
**(A)** Schematic of the TIRF-M assay, proteins and RNAs used. In brief, microtubules are polymerized in vitro from a mixture of purified tubulin, biotin-tubulin and ATTO390-tubulin and stabilized with paclitaxel. These microtubules are then flushed into the assay chamber consisting of a microscopy slide onto which biotinylated, PEG-passivated glass is glued to form a ~5mm wide channel. Microtubules are immobilized by added neutravidin. Then reaction mixtures containing labelled and unlabelled molecules, assay buffer and an oxygen scavenger system are flowed into the chamber. The chamber is sealed, and imaging takes place on an inverted TIRF microscope in a climate chamber set to 30°C. **(B)** Coomassie SDS-gel showing the different full-length APC constructs used in this work. Note, that APC-TMR is a C-terminal SNAP fusion protein labelled with TMR **(C)** Anti-KAP3 western blot and coomassie SDS-gel showing purified KAP3. **(D)** Coomassie SDS-gels showing co-purified heterodimeric and heterotrimeric (KAP3 containing) KIF3AB. **(E)** Anti-KAP3 western blot on gels as shown in (D). **(F)** Simulation of a β2Btubulin-RNA binding curve to APC. The curve was computed assuming a one-site binding model used to fit the MST data in Fig. 1A&B, the Kd measured in Fig. 1A and the APC concentration used in Fig. 3D-F. The red dotted lines indicate that at the β2Btubulin-RNA concentration used in Fig. 3 (2 nM), most APC is saturated with RNA. This shows that the number of transported RNAs measured in Fig. 3 indicate the maximum capacity of the transport system. **(G)** Individual tracks of microtubule lattice diffusing APC-TMR. Each colour represents a different track. The tracks are computed by the Trackmate plugin for Fiji using a minimal displacement cut-off of >1.1 μm to avoid inclusion of unspecific, short binding events into tracks. This way, the entire field of view of entire TIRF-M movies can be included in the analysis to avoid selection-bias. The plot shown was generated using the msdanalyzer package for MATBLAB. **(H)** Individual tracks of transported β2Btub_wt_ RNAs are shown. Here a displacement cut-off of > 4 μm was used to exclude microtubule-lattice diffusing complexes. The track colours in (G+H) represent track IDs. **(I)** The averaged intensity of 12 individual tracks of transported Alexa647-β2Btub_wt_ (2 nM) from TIRF-M assays containing 750 pM KIF3ABK and 80 pM APC is shown. **(J)** The averaged intensity of 12 individual tracks of transported APC-TMR from TIRF-M assays containing 750 pM KIF3ABK, 80 pM APC-TMR and 2 nM TMR-β2Btub_wt_ is shown. The grey lines in I + J are linear regression fits to the averaged intensities. Errors are SD in I + J. Both plots demonstrate that over a time course of 30 seconds, transported RNA and APC do not bleach or significantly loose or gain fluorescence intensity. This shows that no bleaching occurs in this time interval and that the transport complexes on average are not losing or picking up molecules of RNA or APC.

**Fig. S2.**
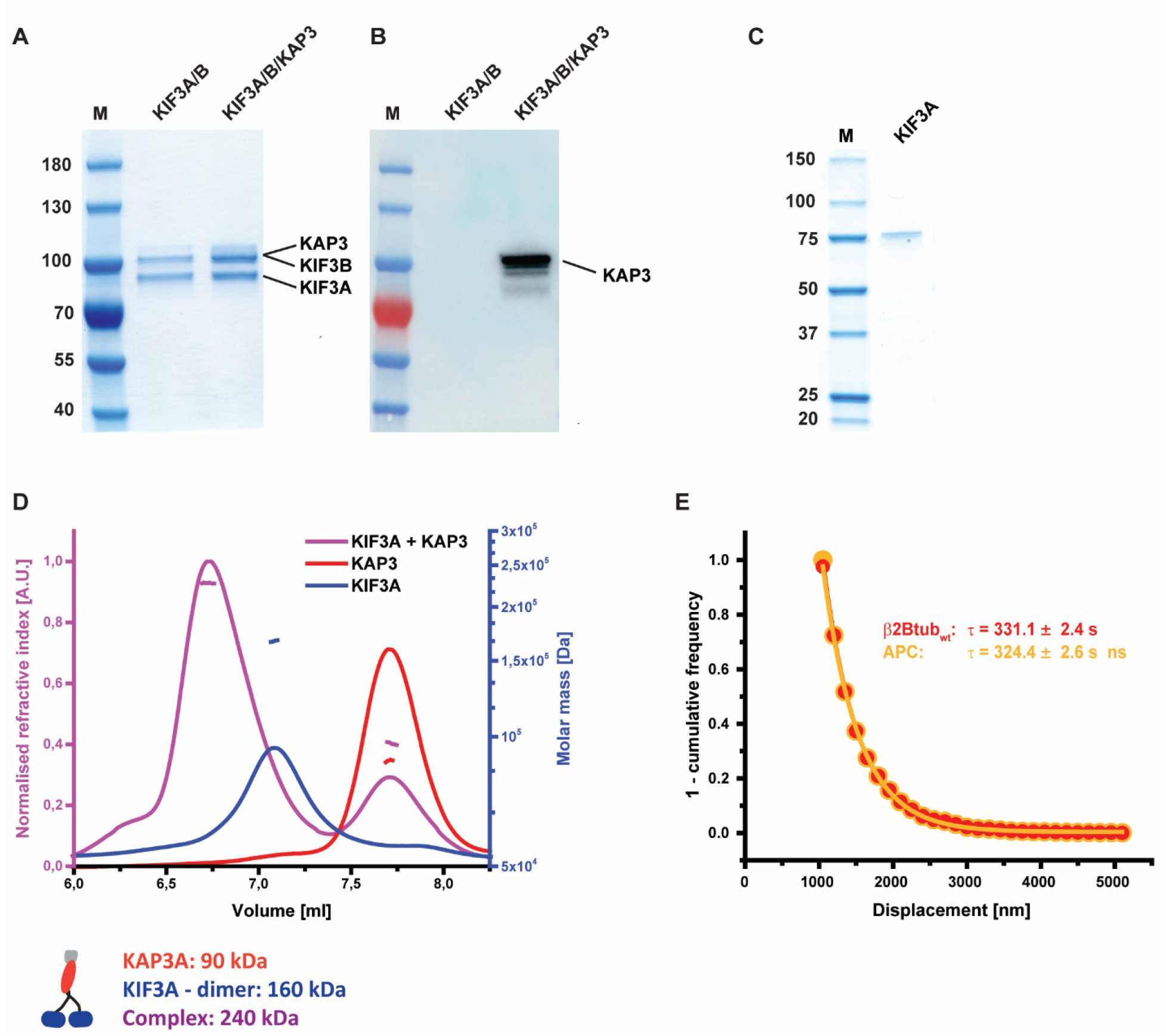
**(A)** Coomassie SDS-gels and western blots of unlabelled, heterodimeric and heterotrimeric kinesin-2. **(B)** Anti-KAP3 western blot of an SDS-PAGE as shown in (A) showing that the purified kinesin contains KAP3. **(C)** Coomassie SDS-gel of KIF3A purified from E. coli. **(D)** Results of Size-Exclusion-Coupled Multi-Angle Light Scattering (SEC-MALS) experiments. The three curves show the elution volume and measured molar mass of KAP3 alone, KIF3A alone and a sample in which KAP3 and KIF3A where combined. The experiments show that KIF3A forms a dimer, which incorporates one KAP3. **(E)** 1-cumulative frequency plot of APC-TMR and Alexa647-β2Btub_wt_ RNPs diffusing on the microtubule lattice. Displacement is defined as the distance between the start point of a track and the end point of a track. The plot demonstrates that diffusive APC-RNPs never show a displacement of more than 4 μm. Hence, we use this value to discriminate diffusive and processive tracks when KIF3ABK is added.

**Fig. S3.**
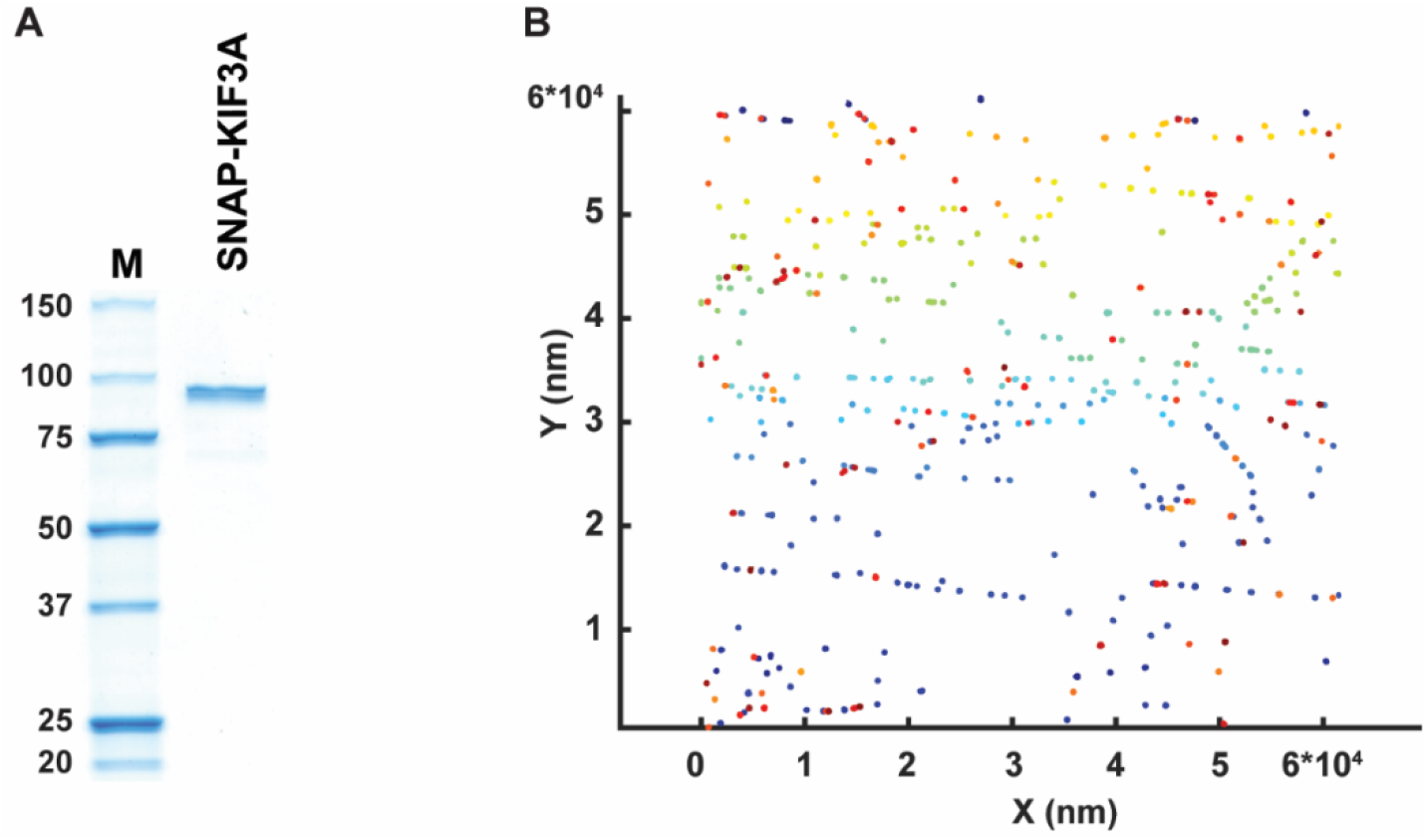
**(A)** Coomassie-stained SDS-gel of KIF3A-SNAP purified from E. coli. **(B)** Individual tracks of microtubule lattice bound Alexa647-KIF3A. Each colour represents a different track. The tracks are computed by the Trackmate plugin for Fiji using a minimal track duration of 3 s to avoid inclusion of unspecific, short binding events into tracks. The plot shown was generated using the msdanalyzer package for MATBLAB. The track colours in represent track number.

**Table S1.**
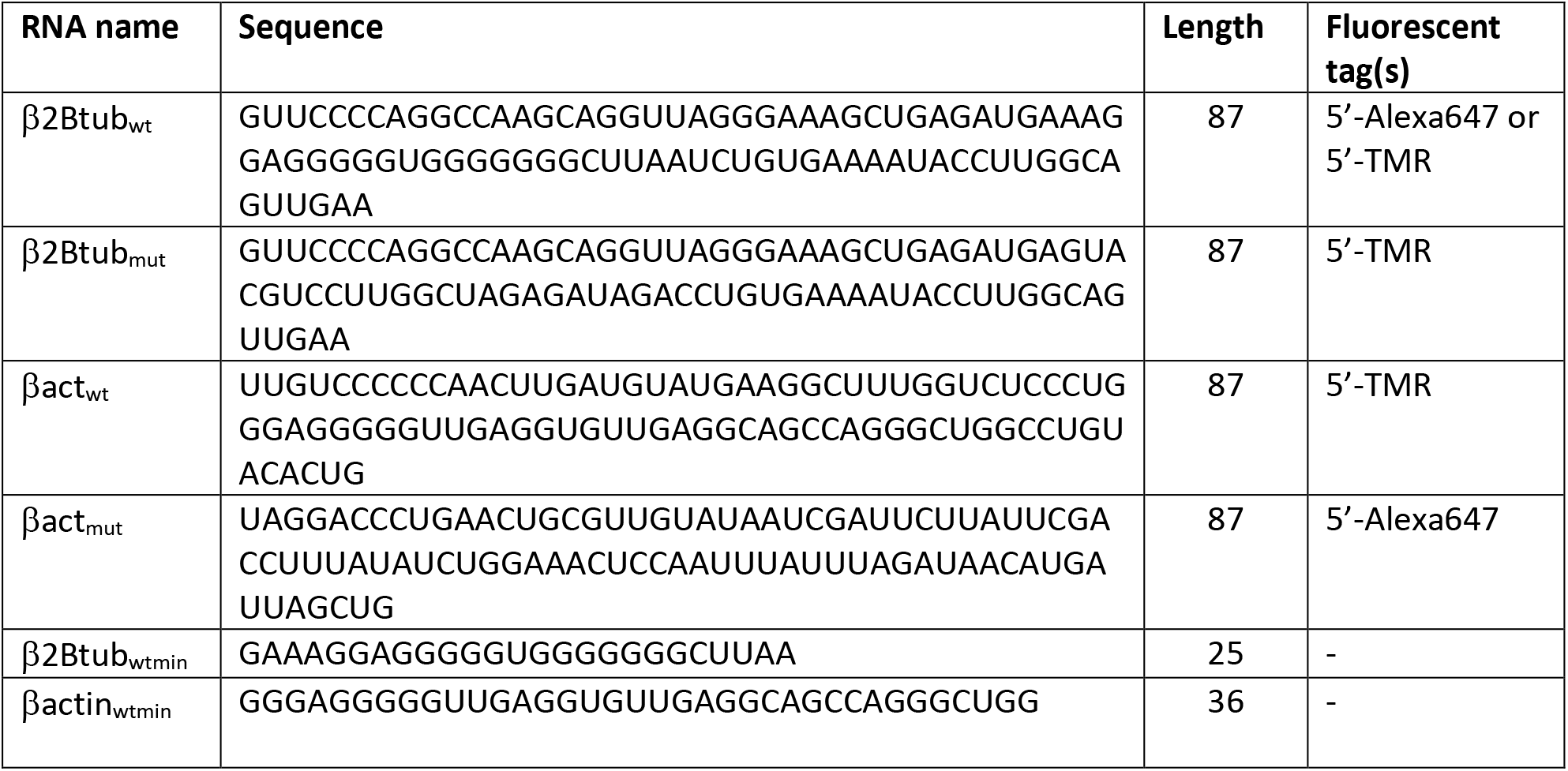
Sequence of RNA-fragments used

**Table S2.** Labelling ratios (in %) of proteins and RNAs used in this study.

|  |  |
| --- | --- |
| Alexa647-KIF3A | 71 |
| Alexa647-KIF3A/B/KAP3 | 73 |
| APC-TMR | 82 |
| Alexa647-β2Btub<sub>wt</sub> | 68 |
| TMR-β2Btub<sub>mut</sub> | 93 |
| TMR-βact<sub>wt</sub> | 92 |
| Alexa647-βact<sub>mut</sub> | 99 |

**Movie S1**

**The kinesin-2 KIF3A/B/KAP3 and APC transport an axonal mRNA.** Dual-colour movie showing the processive and diffusive movement of Alexa-647-labelled β2B-tubulin-RNA (red) on ATTO390-labelled microtubules (grey) in the presence of APC and the heterotrimeric kinesin-2 KIF3A/B/KAP3. The movie was recorded with 2 fps and plays at 50 fps.

**Movie S2**

**Co-transport of APC-β2B-tubulin RNA complexes.** Triple-colour movie showing co-directed movement and co-diffusion of APC-TMR (yellow) β2B-tubulin-RNA (red) on ATTO390-labelled microtubules (grey) in the presence of the heterotrimeric kinesin-2 KIF3A/B/KAP3. Top: APC-TMR on microtubules. Middle: β2B-tubulin-RNA on microtubules. Bottom: Overlay of APC-TMR and β2B-tubulin-RNA with 5-pixel shift in y-dimension. Images were smoothed to assist visualization. The movie was recorded with 4 fps and plays at 50 fps.

**Movie S3**

**APC binds and diffuses on the microtubule lattice in the absence of kinesin-2.** Dual-colour movie showing diffusion of APC-TMR dimers (yellow) on microtubules (grey) in the absence of motor proteins. The movie was recorded with 4 fps and plays at 50 fps.

**Movie S4**

**APC-β2B-tubulin RNA complexes diffuse on the microtubule lattice.** Dual-colour movie showing co-diffusion of APC-TMR (yellow) β2B-tubulin-RNA (red) in the absence of motor proteins. Channels of APC-TMR and β2B-tubulin-RNA are merged with a 5-pixel shift in y-dimension. The movie was recorded with 4 fps and plays at 50 fps. Scale bar 2 μm.

**Movie S5**

**APC recruits and activates the heterotrimeric kinesin-2 KIF3A/B/KAP3.** Top: Alexa647-KIF3A/B/KAP3 (red) on ATTO390-MTs (grey) in the presence of APC. Bottom: Alexa647-KIF3A/B/KAP3 (red) on ATTO390-microtubules (grey) in the absence of APC. The movies were recorded with 4 fps and play at 50 fps.

**Movie S6**

**Single-particle tracking of transported β2B-tubulin-RNA**. Top: Alexa-647-labelled β2B-tubulin-RNA (red) on ATTO390-labelled microtubules (grey) in the presence of APC and KIF3A/B/KAP3. Images were smoothed to assist visualization. Bottom: Same movie as above processed with TrackMate. Circles indicate tracked RNA particles with colour coded total fluorescence intensity. Moving lines indicate local tracks of individual tracked RNAs. Each colour represents a different track. The movie was recorded with 2 fps and plays at 25 fps.

**Movie S7**

**Quadruple-colour movie illustrating the selectivity of the reconstituted mRNA transport system.** Alexa647-β2Btubulin_wt_ (red, wildtype) but not TMR-β2Btubulin_mut_ (cyan, mutated) associates with APC-GFP (green, APC) and KIF3A/B/KAP3 for processive movement along ATTO390-MTs (grey, MTs). Images were smoothed to assist visualization. The movie was recorded with 2 fps and plays at 50 fps.

**Movie S8**

**The APC-KIF3A/B/KAP3 mRNA transport system selectively transports β-actin RNA.** Triple colour movie of a TIRF-M assay containing paclitaxel-stabilized microtubules (grey), TMR-β-actin_wt_ (yellow) and Alexa647-β-actin_mut_ (cyan) as well as APC and KIF3A/B/KAP3. The movie was recorded with 2 fps and plays at 50 fps.

**Movie S9**

**β-actin and β2B-tubulin RNAs are transported in individual packages by the APC-KIF3A/B/KAP3 complex.** TIRF-M assay containing paclitaxel-stabilized microtubules (grey), Alexa647-β2Btubulin-RNA (red) and TMR-βactin-RNA (yellow) as well as APC and KIF3A/B/KAP3. The movie was recorded with 2 fps and plays at 50 fps.

## Notes

#### Summary of Updates

We have updated the article layout to improve readability.

